# Detecting potential reference list manipulation within a citation network

**DOI:** 10.1101/2020.08.12.248369

**Authors:** Jonathan D. Wren, Constantin Georgescu

## Abstract

Although citations are used as a quantifiable, objective metric of academic influence, cases have been documented whereby references were added to a paper solely to inflate the perceived influence of a body of research. This reference list manipulation (RLM) could take place during the peer-review process (e.g., coercive citation from editors or reviewers), or prior to it (e.g., a quid-pro-quo between authors). Surveys have estimated how many people may have been affected by coercive RLM at one time or another, but it is not known how many authors engage in RLM, nor to what degree. Examining a subset of active, highly published authors (n=20,803) in PubMed, we find the frequency of non-self citations (NSC) to one author coming from one paper approximates Zipf’s law. We propose the Gini Index as a simple means of quantifying skew in this distribution and test it against a series of “red flag” metrics that are expected to result from RLM attempts. We estimate between 81 (FDR <0.05) and 231 (FDR<0.10) authors are outliers on the curve, suggestive of chronic, repeated RLM. Based upon the distribution, we estimate approximately 3,284 (16%) of all authors may have engaged in RLM to some degree, possibly opportunistically. Finally, we find authors who use 18% or more of their references for self-citation are significantly more likely to have NSC Gini distortions, suggesting their desire to see their work cited carries over into their peer-review activity.

## Introduction

Quantitative metrics that reflect the potential impact of a researcher’s work or influence of a journal’s papers are highly preferred over subjective metrics, and citations are typically the most influential. For authors, being well-cited can potentially correlate with tangible rewards such as promotions, tenure, and awards, as well as intangible things such as professional and/or societal respect. For journals, citations correlate with quality of future submissions, prestige of being on the editorial board, and potential revenue. Insofar as citations are linked to rewards, individual entities (e.g., researchers, journals) become incentivized to increase the number of citations to their work. Implicit in this is the assumption that citations, particularly citations that come from others, reflect the impact their work has had on their peers. Gatekeepers in the peer-review system, however, have the opportunity and ability to influence references included in a paper. Similarly, authors may feel motivated to include some references not intended to support key points, but to pay homage to some valued entity (e.g., former advisor, departmental chair, colleague, etc). Even outside the peer-review system, there may be ways to manipulate citations, such as by creating documents with citations to your target to be indexed by Google Scholar [1]. Surveys have attempted to estimate different aspects of reference list manipulation, such as how many authors have been affected by coercive citation practices [2], but it’s not known how many researchers have successfully manipulated reference lists in some way for papers they did not author, and how common this may be among active researchers. Furthermore, it would be of interest to know how many researchers, if any, are highly active and persistent in their attempts at reference list manipulation.

The primary purpose of citations is to support key points and link to relevant research. If citations are included in a paper solely to increase someone’s perceived influence, this is deceptive at best. If an editor or reviewer were to use their trusted position within the peer-review process to increase the perceived influence of their own work or of a body of work they have a vested interest in, this is a conflict of interest and unethical behavior. When peer-review gatekeepers request *unnecessary* citations to their own work or journal, this has been referred to as coercive self-citation [3, 4]. For journals, the motivation would likely be to improve their standing (e.g., impact factor), for which the manipulator could obtain either direct or indirect benefits [5]. Authors including unmerited citations (to work other than their own) could be the result of a favor or a *quid pro quo* arrangement between authors which, when recurring, are referred to as “citation rings” [6, 7]. Editors influencing unmerited citations may be a result of coercion but, depending upon the manuscript handling process, it’s also possible they could simply insert citations to their own work after a paper is accepted without the knowledge of the authors. Unmerited journal self-citation can be achieved either by encouraging authors to cite more papers from their journal [4] or simply by publishing content with self-citation. When journals unified by a common interest (e.g., same parent company or editor) cite each other, this is called “citation stacking” [8]. For a more thorough overview, Ioannidis has a written a very helpful commentary discussing the many different types of self-citation, depending on where they originate and how they manifest [9].

The goal of this study is to develop and test a method to detect patterns of reference list manipulation (RLM). More specifically, we are interested in estimating how many authors have engaged in RLM for the purpose of benefitting themselves, non-transparently, within the peer-review system. We thus propose the term “citation hacking” may be a more descriptive term for this. Hacking, as a phenomenon, involves unauthorized, non-transparent access to a system that is otherwise presumed secure and/or trustworthy, such as peer-review and publishing, to perform some action that benefits the hacker. In this case, the hacker’s goal would be to increase the perceived influence of a body of research. Although the most likely beneficiary would be the citation hacker, either directly (to their own research) or indirectly (to journals they are founders of or editors for), it’s conceivable that the intended beneficiary could also be a group of people united by some common factor (e.g., university, department, co-investigators on a grant, etc), or even a favored theory that is in dispute. Thus, the term “citation hacking” encompasses the use of non-transparent means to achieve an end goal (to increase perceived influence) and the compromised system (publication & peer-review), but neither the means used nor entity responsible (editor, author or reviewer). By this definition, self-citations (SC), because they are transparent, are not citation hacking.

There are three important issues that motivate this study. First, although surveys have approximated how many authors have been victims of coercive citation, it is not known how many researchers may have engaged in citation hacking and to what degree (e.g., many to some extent, a few to a large extent, or both). Related to this point, it is also of interest to identify potential risk factors that might predict future behavior. Second, as we noted in a previous publication [10], even after citation hackers are discovered, because of privacy concerns and a highly decentralized publishing system, there is no effective mechanism to share information. Even though there are checks-and-balances built into the peer-review system (e.g., editors screening reviewer reports), from their standpoint, they are often only witnesses to a single incident and reluctant to raise concerns on that basis alone [11]. Furthermore, there is little incentive to publicly disclose citation hacking events once uncovered. In fact, there are potential disincentives such as reputational embarrassment and potential litigation risks if naming offenders. Given the large, decentralized ecosystem of journals, citation hackers can potentially continue unabated as reviewers and/or editors simply because the number of people aware of their behavior within the ecosystem will be a tiny fraction of the whole. Third, under the presumption that many authors may not describe their requests for self-citation as unmerited, even when excessive, it is important to be able to put both their individual requests and observed patterns over time within the context of a peer group (in this case, highly published authors).

In summary, we know citation hacking to be a problem, but without understanding the scope of the problem, we have no way to prioritize prevention efforts. And to disincentivize the behavior, we need a method to detect it that is fairly simple to use and understand (like the H-index) and can quantify deviations from the norm.

## Background

A 2008 survey of 283 authors found 22.7% of them reported “*a reviewer had required them to include unnecessary references to his/her publication(s)*” [12]. Although this survey was relatively limited in its scale and scope, and the key word “unnecessary” does not necessarily imply that the references were inappropriate or excessive, but it does suggest two things: First, by virtue of the number and nature of the citation requests, the author believes they were able to infer the reviewer’s identity. Second, requests for additional citations that, in the author’s view, are not merited may be fairly common in scientific peer-review. It is not clear, though, what fraction of these requests might be characterized as ego-driven (e.g., demanding recognition that they have contributed in a related area) versus what fraction might be a deliberate attempt to increase the reviewer’s perceived influence. Similarly, another survey found more than 20% of respondents had experienced coercion from a journal editor [4].

Another study reported that 29% of the references that reviewers requested the authors add during peer-review were to the reviewer’s own work [3]. The study, however, did not ascertain what fraction was perceived as unmerited. However, in reviews recommending acceptance/revision, more than twice as many reviewer self-citation requests were found than in those recommending rejection (reported p<0.001), whereas the number of requested citations to the work of others did not significantly differ. Interestingly, a related study found that requested self-citation frequency did not differ between blinded and open peer-review [13], suggesting author awareness of a reviewer’s identity is not a significant disincentive. Instances of various types of citation hacking have been documented, but are hard to detect in general [9, 10]. This is the first study to attempt a literature-wide, quantitative estimate of how many authors may have engaged in citation hacking and how many chronic offenders may be currently active. To do this, we analyzed references from recent papers (mostly from the past 10 years) that cited a subset of currently active, highly published authors in MEDLINE.

## Methods

### Obtaining citation data

PubMed 2020 records were downloaded in XML format on May 25, 2020 from NCBI (ftp://ftp.ncbi.nlm.nih.gov/pubmed/). For each paper, we extracted its PubMed ID (PMID), all author names, name of the journal it was published in, the journal’s ISSN, and the PMIDs of each paper within the reference list, when given. Only references that contain a PMID are included in the citation network (i.e., a paper may have more total references than the ones extracted).

**Figure 1** shows the distribution, by year, of citations to papers and references from papers. Since references are extracted from papers deposited in PubMed Central (PMC), they are heavily biased towards more recent papers, although citations to papers extends much further back. Within this dataset there were 31,029,833 unique PMIDs, 6,003,225 (18%) of which contained at least 1 reference, and 16,338,882 (53%) of which received at least one citation. A total of 172,528,049 PMID-PMID citation links were identified.

**Figure 1:**
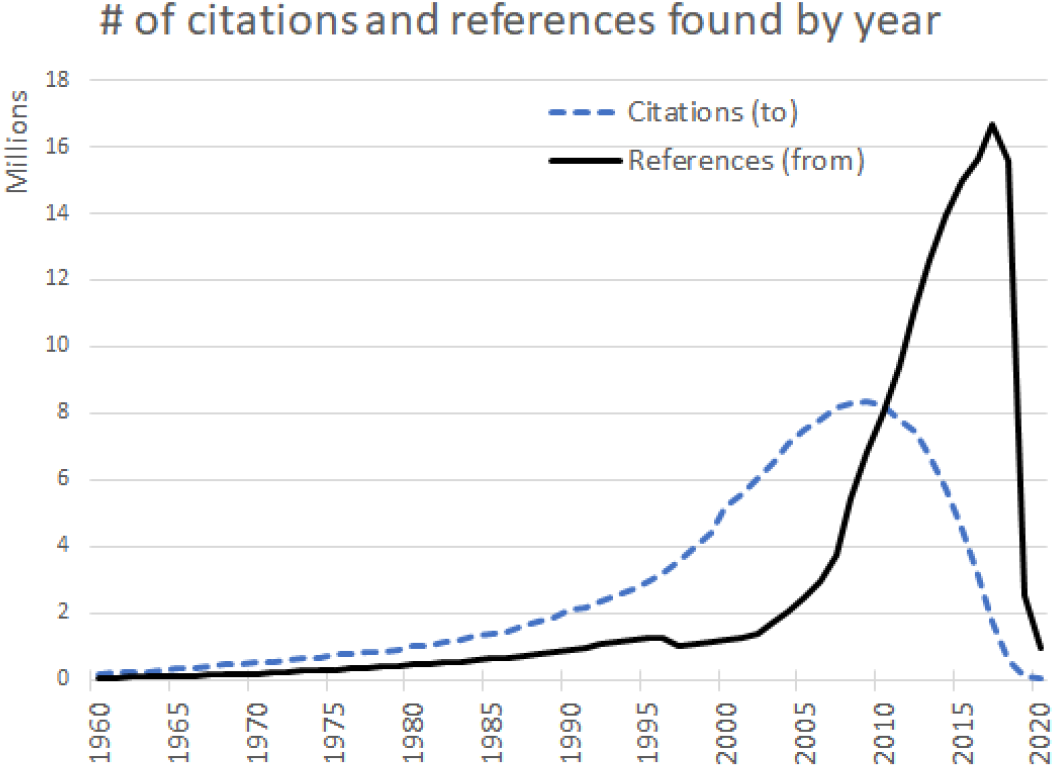
The total number of references found in papers by year (black line). The dashed blue line shows the years in which the cited publications were published. There is an apparent lag time to entry of citation data, so citations to the most recent papers are sparse, which appears as a sharp drop-off.

References in the XML records tend to be given in order of their appearance within the full text, even if the journal’s publication reference format is alphabetical by author name. This enables analysis of over-represented author names within blocks of contiguous references, but we note that, upon comparing several extracted lists with the PDF of the publication, there were some discrepancies in the ordering, suggesting the mapping is approximate, not exact. Thus, it is problematic to accurately quantify, using this data, the largest contiguous block of citations to one author, but less problematic to estimate how many smaller contiguous blocks exist.

### Subsetting authors for analysis

We restricted our author list to include only authors who have published recently (most recent publication within the citation network no older than 2017) and to those who authored or co-authored at least 100 papers within the citation network (i.e., 100 papers with at least one citation to or from the paper). We denoted authors in the first and last positions as “anchor” authors, because they are considered to have had a disproportionate influence on the content of the paper [14]. When we refer to “authorship”, this includes co-authorship, and is defined simply as the presence of an author’s name on the authorship byline. Accented letters were converted (e.g., “Peña” => “Pena”) to reduce potential inconsistency in transcription (e.g., by a co-author when writing, a journal when formatting, or PubMed when indexing).

Because our analysis is author-centric, in the absence of a widespread unique author identifier within the metadata (e.g., ORCID), it was necessary for us to attempt to reduce the level of author name ambiguity. We did this by restricting analysis to authors with names that included at least one middle initial or, if not, then had either two first or last names (as judged by the presence of either a hyphen or space within the first/last name). Also, we required their middle plus first name to be at least 4 characters in length. One downside to this limitation is that it will underestimate the total number of citations to authors in proportion to the number of inconsistencies in their full name (e.g., “Smith, JA”, “Smith, John Abrams”, and “Smith, John A” would be counted as different people even if they refer to the same person), or if the author underwent a name change at some point (e.g., due to marriage). Also, because SC rates are higher than NSC rates, there is a risk that authors with high SC rates that are also highly inconsistent in the spelling or structure of their names will be detected as having highly distorted NSC. A total of 20,803 authors fit all these criteria. Ambiguity of author names, by these criteria, appears to be more of a problem prior to 2002 than after, perhaps because of a shift from recording author first/last names as initials to their full-form. It is difficult to estimate what fraction of the whole this subset represents because, although there were over 12 million unique author name strings identified, some authors will share a name. Furthermore, ~46% only occur once, some of which will be attributable to name variations (including spelling errors), and others to people who have only published once.

### Normalizing metrics

Many of the red flag metrics scale with an author’s total number of non-self citations (NSC). To more effectively remove the influence of total NSC, we subtracted out the effect of total NSC by modeling the relationship between the two variables using robust linear regression, in log-log space. Log transformations were determined using the box-cox power transform to be optimal for meeting standard regression model assumption requirements, including linearity, normality and homoscedasticity. Exponentials of the regression residuals return the observed vs expected ratios, and are shown in graphs as normalized values. In raw form, in log-log space, the regression residuals are approximately normally distributed, providing suitable inputs for Factor Analysis.

## Results

### The number of non-self-citations (NSC) to one author from one paper approximates a Zipfian distribution

Citation hacking, by definition, takes place on the level of the individual paper. The frequency of citations, within one published paper, to any of a single author’s entire body of published papers, is approximately power-law distributed or Zipfian. For our data, we find linear projection (Ordinary Least Squares) of NSC frequency on NSC number, on a log-log scale explains more than 90% of variability (R^2^ >0.9) for more than 95% of the authors. **Figure 2** shows this linear log-log relationship, characteristic of a Zipfian distribution, is roughly valid overall, for our author subset. Self-citations (SC) did not follow a Zipfian distribution, but this was expected since they are governed by different mechanisms (i.e., an author’s preference as opposed to external awareness/interest).

**Figure 2:**
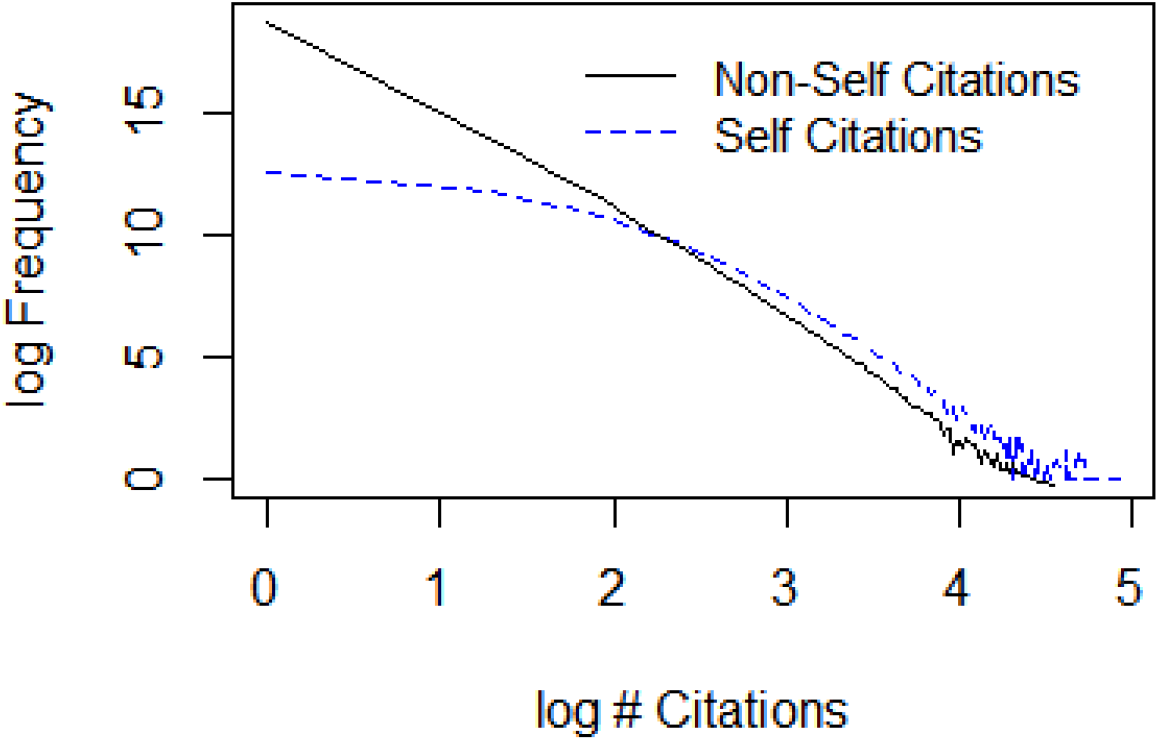
Cumulative # of times (y-axis) one paper cited one author exactly *n* times (x-axis) (NSC=non-self citations, SC=self-citations), plotted as the natural logarithm of the number.

Zipfian distributions are known to arise in a variety of natural systems and are thought to be governed by laws of preferential attachment [15, 16]. An important implication of Zipf’s law to this study is that, because the trend is linear in log-space, future values in the series can be approximated using the initial or early values in the series. Thus, the frequency by which any author has received *n* citations from a paper should be proportional to the frequency by which they receive *n*+1, *n*+2, etc. This frames the problem overall by defining statistical expectations regarding what is “normal” and enabling null hypothesis testing for observed frequency distributions. It does not mean that authors with skewed distributions have necessarily engaged in RLM, but it does mean that all authors who do engage in RLM will have skewed distributions in proportion to their activity. The more references requested per attempt and the more frequent the requests, the more severe the skew.

### Identifying “red flags” suggestive of citation hacking

Because there is no gold-standard or ground truth to evaluate how well a metric reflects citation hacking activity, and because citation hackers may have different strategies and/or opportunities to influence reference lists, we examined several patterns that are suggestive, but not independently conclusive, of citation hacking (“red flags”). Because of the highly sensitive nature of implicating an author as one who has engaged in RLM, it’s critical to have interpretable metrics that provide a basis for further investigation if needed (**Table 1**).

**Table 1:** Summary of red flags used to identify patterns of behavior that are suggestive of RLM.

| Red Flag # | Base metric | Abbr. | For each author, an unusually high frequency of: |
| --- | --- | --- | --- |
| 1 | NSC | Blocks | Consecutive citations to them within papers |
| 2 | NSC | NSCI | Papers with multiple citations coming from them |
| 3 | NSC | 17+ | Extreme citation events |
| 4 | NSC | MCJI | Multiple citations in papers published in a specific journal |
| 5 | SC | %SC | Self-citation, as judged by % of reference list dedicated to it |

Each flag is motivated by economic considerations: If someone wants to increase their perceived influence via increased citation of their work, but their supply of opportunities is limited, then their incentive is to maximize the number of references added per opportunity available. This is mitigated by other factors such as author compliance and editorial intervention, but also by some expectation the hackers have regarding potential costs associated with their behavior being called into question.

### The Gini Index as a proposed metric for quantifying skew in a frequency distribution

The Gini Index is a well-known statistical measure of dispersion and inequality, especially popular in economics to quantify inequalities of income distribution that ranges from 0 (perfect equality) to 1 (perfect inequality). In this case, it will measure how skewed the NSC frequency distribution is from individual papers to an author. If each data point contributes approximately equally, then the Gini will be close to 0. The more an author receives large numbers of citations from individual papers relative to the times they receive fewer citations, then the closer to 1 their Gini will be. One advantage of the Gini is that, as a scale-independent relative measure, it does not require normalization. The distribution of Gini index values for all authors is shown in **Figure 3**.

**Figure 3:**
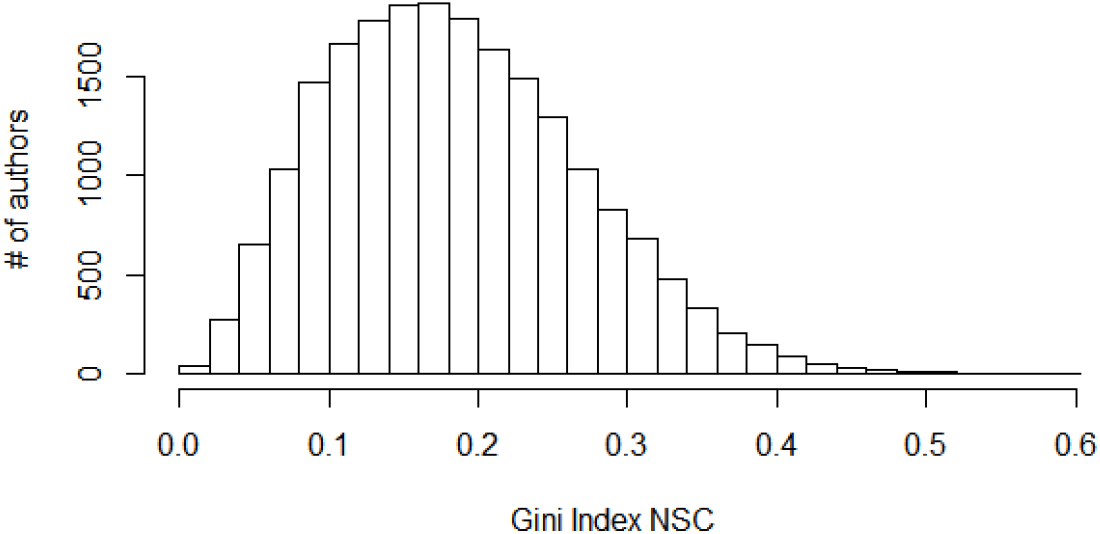
Distribution of Gini Index values (mean=0.185 ± 0.084), quantifying how skewed the single-paper NSC frequency distribution is for each author. The higher the Gini, the more often an author had unusually large numbers of NSC to their work coming from single papers, relative to their overall distribution.

To examine how sensitive the Gini is to outliers, we removed the paper with the most NSC for each author, recalculated their Gini and compared the rankings using Spearman’s correlation. We found a high concordance (R=0.996) and that within the top 1% highest Gini’s (n=208), the most an individual author dropped was 32 positions, suggesting their membership at the extreme end was not sensitive to single outlier removal.

We also wanted to examine whether or not the Gini might be influenced by “mega-reviews”. Mega-reviews are papers with an unusually high number of references that attempt to summarize work within a large area of research. Thus, they might be prone to citing individuals in the field more frequently in a single paper. Since there is no standard definition for how many references make a paper a mega-review, we recalculated Ginis after excluding NSC from all papers with >150 references, which encompasses only 0.8% of all papers but 6.3% of all references. Within the top 1%, one author’s Gini fell by a striking 788 positions, but the average drop for the remaining authors was only 1 rank. Further examination of this author shows 128/991 (13%) of the papers with >=1 NSC to them also had >150 references, suggesting 150 may be an insufficiently small cutoff for some fields, and showing that this drastic change in Gini was not attributable to a small number of mega-reviews.

Finally, we estimated how much their Most Citing Author (MCA) affected the Gini of each author in the list, as defined by an H-index-like measure of at least *n* NSC observed in at least *n* papers for each citing author. Within the top 1%, the ranking for two authors dropped substantially (**Figure 4**), but most did not change appreciably. Note that Gini contributions from single outliers and mega-reviews are viewed as potential confounds, but the MCA may or may not be innocuous (i.e., they may simply be admirers or a *quid-pro-quo* might exist). Based on string similarity of each author’s name versus their MCA, we estimate that about 1.4% of the authors have an MCA that is actually a variation on the spelling of their own name. This suggests that mistaking SC for NSC is happening at a fairly low rate.

**Figure 4:**
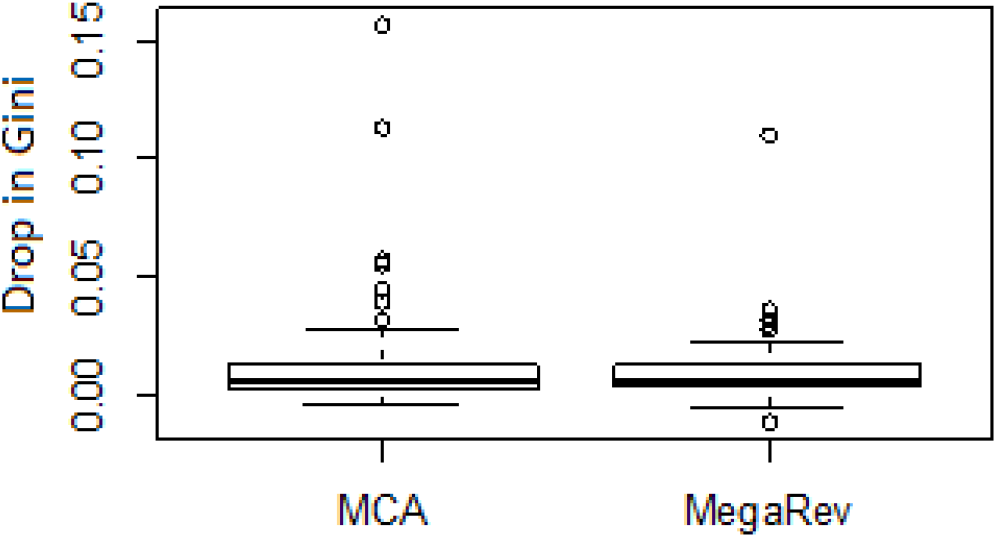
Drop in Gini for authors with the highest 1% NSC Gini Index scores when subtracting their Most Citing Author (MCA) or all mega-reviews (papers with >150 references).

### Red Flag #1: Seeing large and/or frequent blocks of consecutive NSC to an author within a paper

References are expected to support the topic at hand and, although there may be valid reasons for an author-centric series of consecutive references, it is not the norm. Unusually large and/or frequent blocks of contiguous NSC are highly suggestive authors may be accommodating a request from either a reviewer or editor. Not only did we observe this for the coercive reviewer documented in our case report [10], but it makes sense that when authors are adding citations solely to satisfy a reviewer’s concerns, they would generally do it in one or possibly a few blocks of consecutive citations. The alternative would be to try to weave them throughout the text and, particularly for unmerited citations, would be quite difficult to do in a way that appeared natural or logical to the reader. Thus, we expect that a common “fingerprint” left by citation hackers would be large blocks of contiguous citations to them within a paper. This metric lends itself to validation by examining the surrounding context of citation blocks. The more generic the statement (e.g., “other work has been done in this area”) and the larger the block, the less likely the citations were motivated by the topic or necessary to the paper. Considering authors with at least 200 NSC (n=20,712), the ratio of total consecutive NSC (3 minimum) to total NSC shows that such citation blocks are relatively uncommon events in general, with 30% of authors having none at all (**Figure 5**).

**Figure 5:**
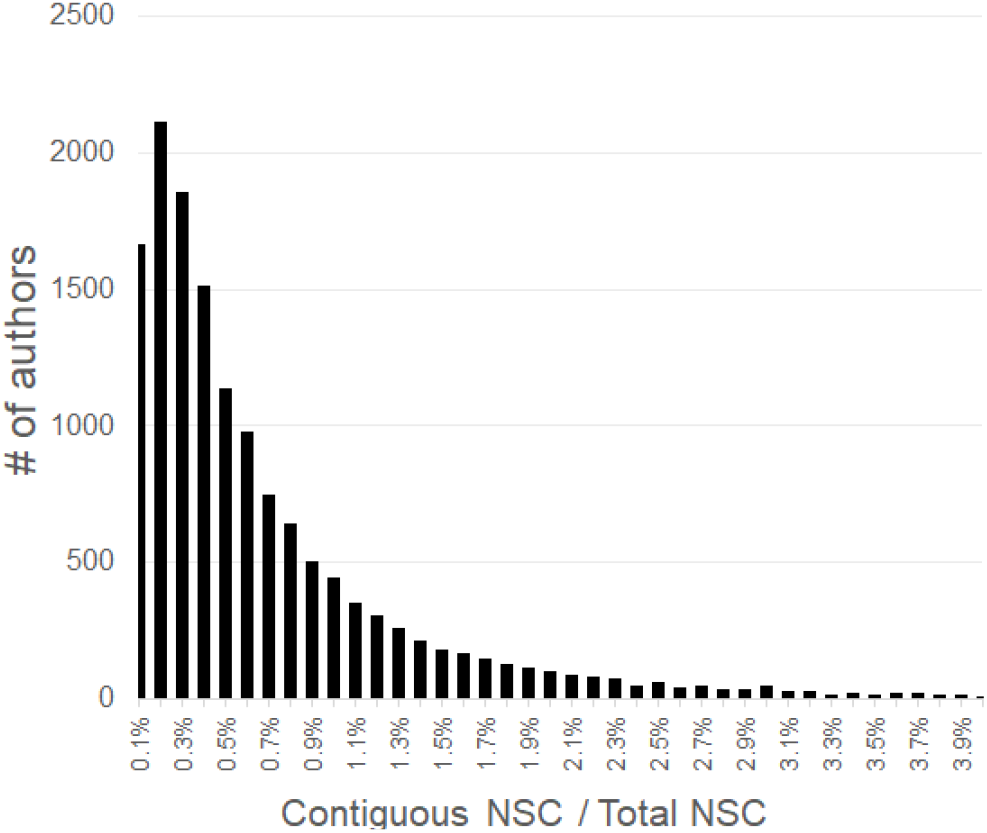
Histogram showing how the percentage of contiguous NSC (minimum of 3) to total NSC is distributed for each author (x-axis truncated at 4%, maximum value = 64.4%).

### Red Flag #2: Multiple papers with an unusually large number of NSC to an author relative to their total NSC

**Supplementary Table S1** shows the probability of observing n citations to one author’s work in a paper, both NSC and SC. For example, although it is rare to see more than 5 references to someone who is not an author of the paper (~1%), it is fairly common to see more than 5 citations to one of the paper’s authors (23%). Thus, conceptually similar to the H-index, we can calculate an NSC Index (NSCI) using the number of times, *n*, that at least *n* NSC came from one paper to a specific author. Because the H-index correlates with the square root of the total number of citations [17], the NSCI is normalized (see methods). **Figure 6** shows the distribution of NSCI values for all authors.

**Figure 6:**
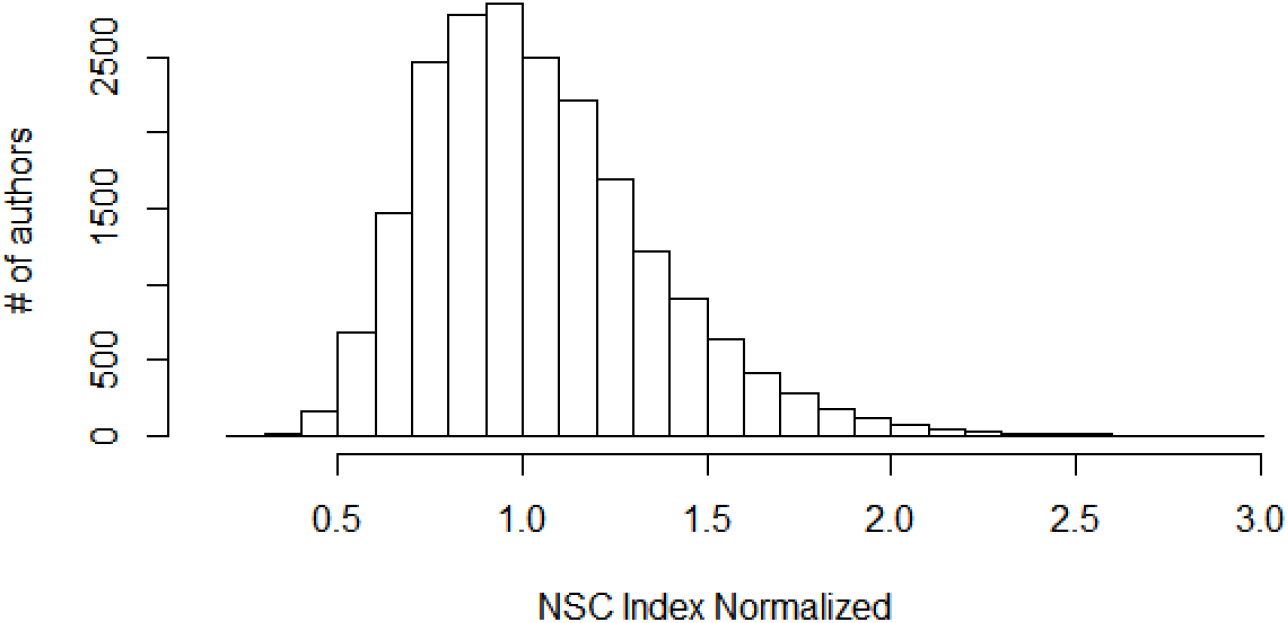
A normalized H-index like measure for NSC coming from individual papers to an author shows a central tendency, but with a skew towards higher values.

### Red Flag #3: Papers that contain an extremely large number of NSC to an author’s body of work

The NSCI, similar to the H-index, seeks to discount the extreme end of the citation curve in favor of a metric that more stably reflects the entire distribution. However, extreme events are not only informative, but reflect how egregious the hacker can be and also represent instances whereby the peer-review system has clearly broken down. For example, if someone coerces the insertion of 49 references to their work, the NSCI could detect this if evenly spread out (7 papers with 7 NSC each), but not if they are all in one paper. Similarly, it seems less concerning to discover an editor did not question or notice a reviewer requesting 7 self-citations in 7 separate reviews spread out over time versus an editor who did not question or notice a reviewer who requested 49 self-citations in one review. Whereas the NSCI prioritizes consistency over egregiousness, the observed to expected ratio of extreme events prioritizes egregiousness over consistency. Note: There may be valid reasons for a paper to contain an extreme numbers of NSC to one author (e.g., honoring a retired or deceased author). However, we expect this to be relatively infrequent for most authors, but much more common among citation hackers.

As **Supplementary Table S1** shows, the odds that 17 or more NSC will come from one paper to one author is approximately 0.025%, or once per 3,942 papers that contain at least one NSC to an author. 17 was chosen simply because it is a high threshold that exceeds the average NSC per author (3,062), and 82% of authors in our subset have never even received 17+ NSC. Within the 12,110 times 17+ NSC to one author came from one paper within our subset, they contained a total of 261,067 citations (avg=22). For authors with at least one 17+ NSC, an expected number of such events is computed by modeling the increase with total NSC using Poisson regression (**Figure 7**).

**Figure 7:**
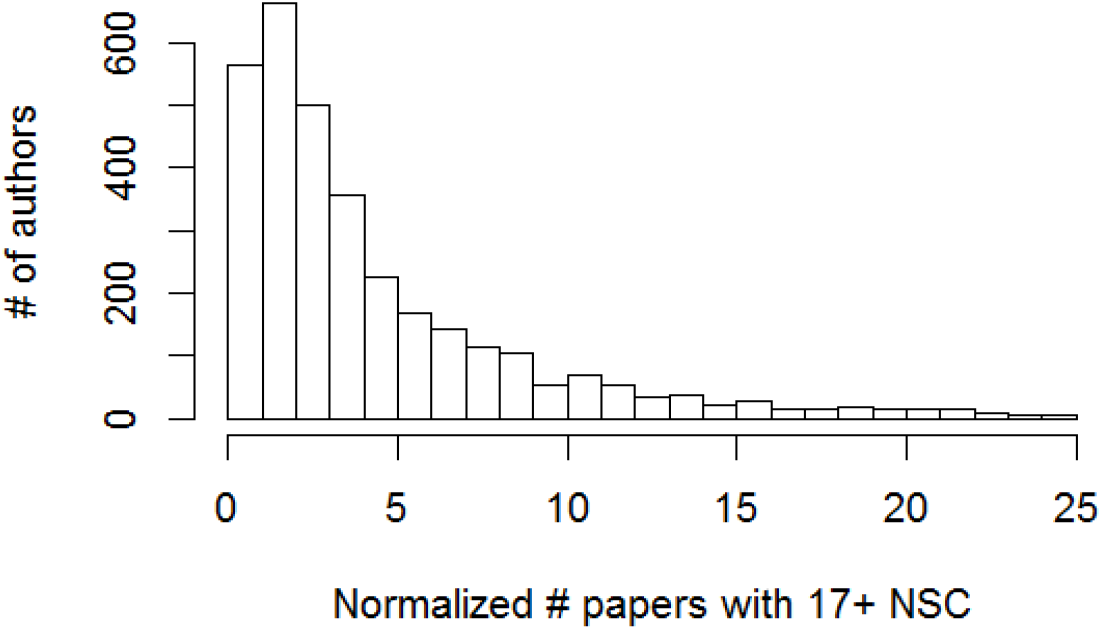
Observed to expected ratio of number of extremely large non-self citation events (17+) to one author from individual papers, controlling for author’s total NSCs.

### Red Flag #4: An unusually large number of NSC to one author coming from papers published within one journal

A high number of NSC to one author from papers published in a specific journal are suggestive of a researcher who may have requested citations to their work in their capacity as handling editor or, possibly, a reviewer frequently used by an editor. First, for each journal publishing a paper with at least one NSC to an author, an H-index like measure, MCJI, is calculated reflecting the largest number of papers, *n*, whereby at least *n* NSC were observed. The journal with the highest *n* is denoted here as the Most Citing Journal (MCJ) for that author. **Figure 8** shows the distribution of MCJI values normalized to author’s total number of NSCs from MCJ journal.

**Figure 8:**
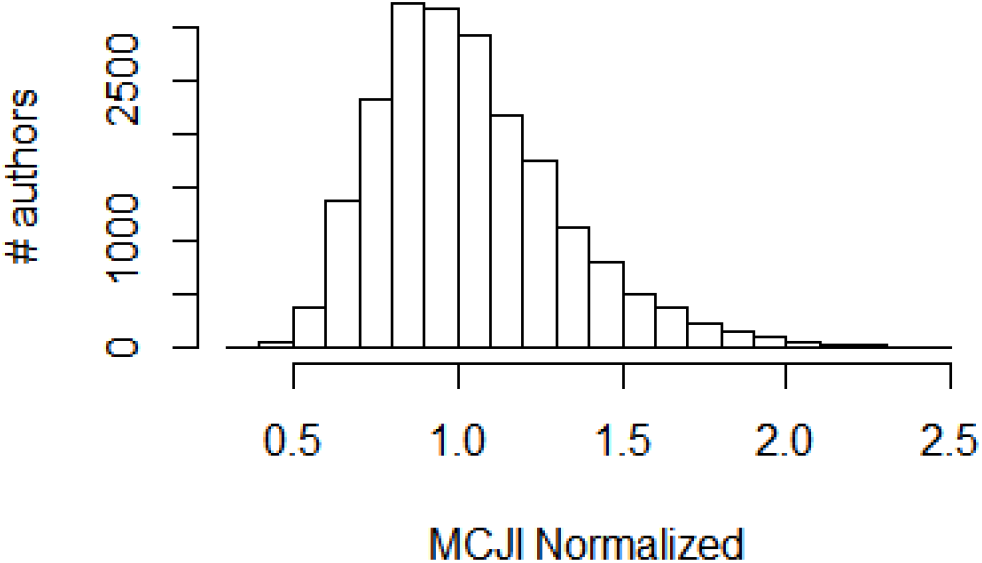
Histogram of normalized MCJI for each author.

Note: Although a high MCJI may be informative, a low MCJI could reflect lack of editorial appointments or reluctance to use that venue for citation hacking. For example, unlike reviewers, editors lack anonymity and their requests would be seen by both authors and reviewers. In fact, a recently published case of an editor using his position to coerce reviewers into citing his papers found he created email pseudonyms so that requests to cite his papers appeared to be coming from a reviewer [18].

### Red Flag #5: Self-citation at the cost of excluding field coverage

Excessive self-citations, as measured by the fraction of space reserved in an author’s reference list for self-citation, are suggestive that an author is not merely attempting to draw attention to their prior work but, more specifically, using the publication opportunity to maximize the total number of citations to their work. SC are generally transparent and can be subtracted from metrics such as the H-index, when desired. But given that SC may or may not be subtracted, for someone who wants to increase the perceived influence of their work, there is no reason to restrict their efforts to only papers they handle during peer-review, particularly when there is no consensus on whether or not excessive self-citation is an ethical breach [19, 20]. Thus, there is potential reward without much risk. **Figure 9** shows the distribution of fractional self-citation per author, for authors with a minimum of 9 “anchor” author (i.e., 1^st^ or last author) papers with at least 100 total references among them, broken down by anchor vs middle.

**Figure 9:**
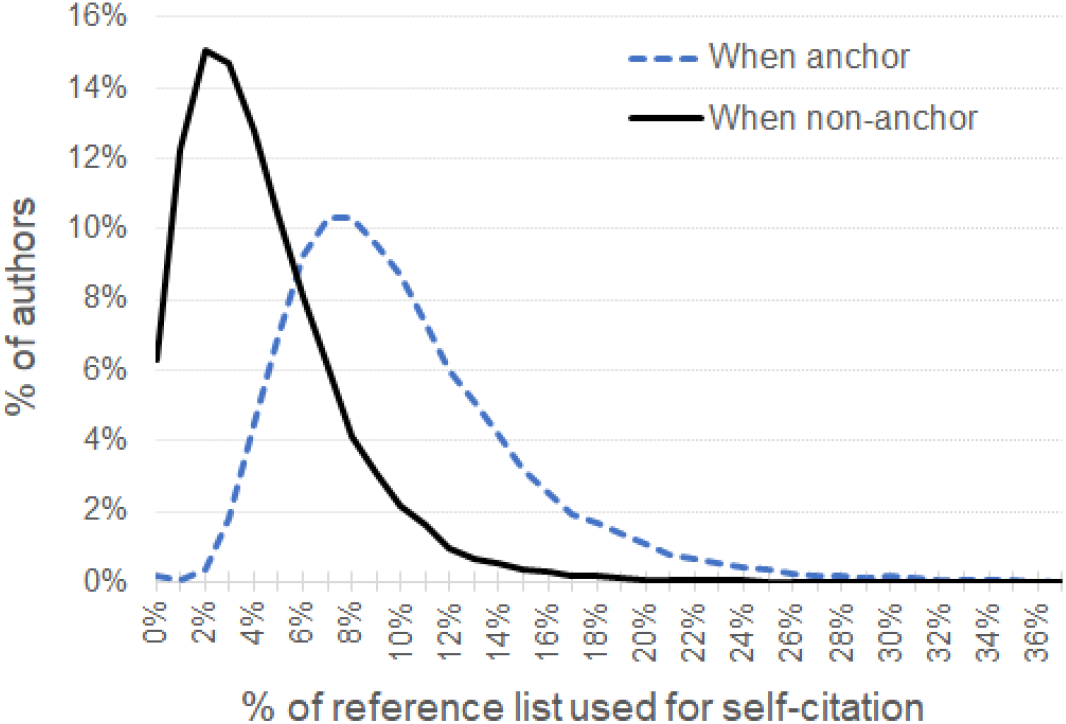
Percent of the reference list used for self-citation within our author subset (x-axis truncated at 36%, maximum value=66%). Self-citations are more common when an author is either the first or last author on the paper’s byline (i.e., “anchor’s” one side of the byline).

### Gini captures 95% of the ranking information from other metrics

Examining the correlation structure of each red flag metric (**Table 2**) shows each one contains information about the others to some degree, but are not so highly correlated that they are redundant. While correlations among the first five NSC-based variables were expected, we were surprised initially to see such a strong correlation between NSC-based metrics and %SC which, in theory, should be completely unrelated.

**Table 2:** Spearman’s rank correlation among “red flags” that are suggestive of citation hacking. Gini=Gini index of NSC distribution; NSCI = Non-Self Citation Index; %SC= average percent of the reference list used for self-citation; Blocks=citations in contiguous blocks (>=3); 17+= Papers with 17 or more NSC to one author; MCJI= Most Citing Journal Index; Factor= aggregate factor analysis score.

|  | Gini | NSCI | Blocks | 17+ | MCJ | %SC | Factor |
| --- | --- | --- | --- | --- | --- | --- | --- |
| Gini | 1 | 0.82 | 0.63 | 0.48 | 0.68 | 0.55 | 0.95 |
| NSCI | 0.82 | 1 | 0.66 | 0.52 | 0.61 | 0.52 | 0.93 |
| Blocks | 0.63 | 0.66 | 1 | 0.43 | 0.47 | 0.51 | 0.75 |
| 17+ | 0.48 | 0.52 | 0.43 | 1 | 0.39 | 0.35 | 0.58 |
| MCJ | 0.68 | 0.61 | 0.47 | 0.39 | 1 | 0.38 | 0.73 |
| %SC | 0.55 | 0.52 | 0.51 | 0.35 | 0.38 | 1 | 0.63 |

Factor analysis, similar to Principal Component Analysis, searches for one or more latent/unobserved variables that best explains joint variation within a group of variables. Here, the first latent factor accounts for 55.4% of the group’s total variation, and the second factor only 11%. Factor scores for each variable suggest how well they reflect the behavior of the group. Notably, Gini has the highest predictive power, explaining 51.3% of the first latent variable. This, plus its relative simplicity and the fact it does not require normalization, makes it a good metric to detect potential citation hacking. **Supplementary Figures 1-7** show plots of each factor versus the others, normalized and non-normalized.

### Estimating levels of chronic and acute citation hacking

Since the Gini coefficient can be seen as the covariance between a variable and its rank [21], and covariance is itself a mean that converges to a normal distribution, others have used Gaussian distributions to assess Gini values statistically [22]. Under the assumption that the distribution of NSC Gini index values is approximately Gaussian, we can estimate two things. First, we can assign a statistical confidence by which we can reject the null hypothesis that an author’s Gini index value is part of the normal distribution. Chronic citation hackers who engage in repeated and/or egregious RLM would be expected to fall well outside the norm. Second, under the additional assumption that authors would rarely ask for removal of references to their work from papers they review or handle as editor, then in a world where RLM did not exist, the left and right-hand sides of the curve should be symmetric. Because we do know, at least from a limited number of reports, RLM has happened, then the extent to which the real-world curve is right-shifted relative to the ideal curve provides us with a quantitative estimate of the difference.

The black line in **Figure 10** shows the distribution of Gini values for all authors, whereas the red dotted line shows a Gaussian distribution, with a mean and deviance that best fit the left-hand side of the curve. We then compute for each Gini number the p-value by which we can reject the null hypothesis that it is part of the reference distribution, correcting for false discovery rate (FDR). Authors with the lowest FDR values will correspond to those who have an abnormally large NSC Gini index and by which the null hypothesis that such patterns are normal can be confidently rejected. The full list of authors and their FDR p-values is in **Supplementary Table 2**, which is available upon request. There are 81 authors (0.4%) with FDR<0.05 and 231 with FDR <0.10 (1.1%). Summing 1-FDR across the entire set, provides us with an estimate that 3,284 (16%) of the authors in our subset have higher Gini values than expected (**Figure 10**, grey area). Note that this is a population-level statement, not a threshold to evaluate individual Gini scores. It suggests that about 16% of authors may have engaged to some degree, on one or more occasions, in RLM.

**Figure 10:**
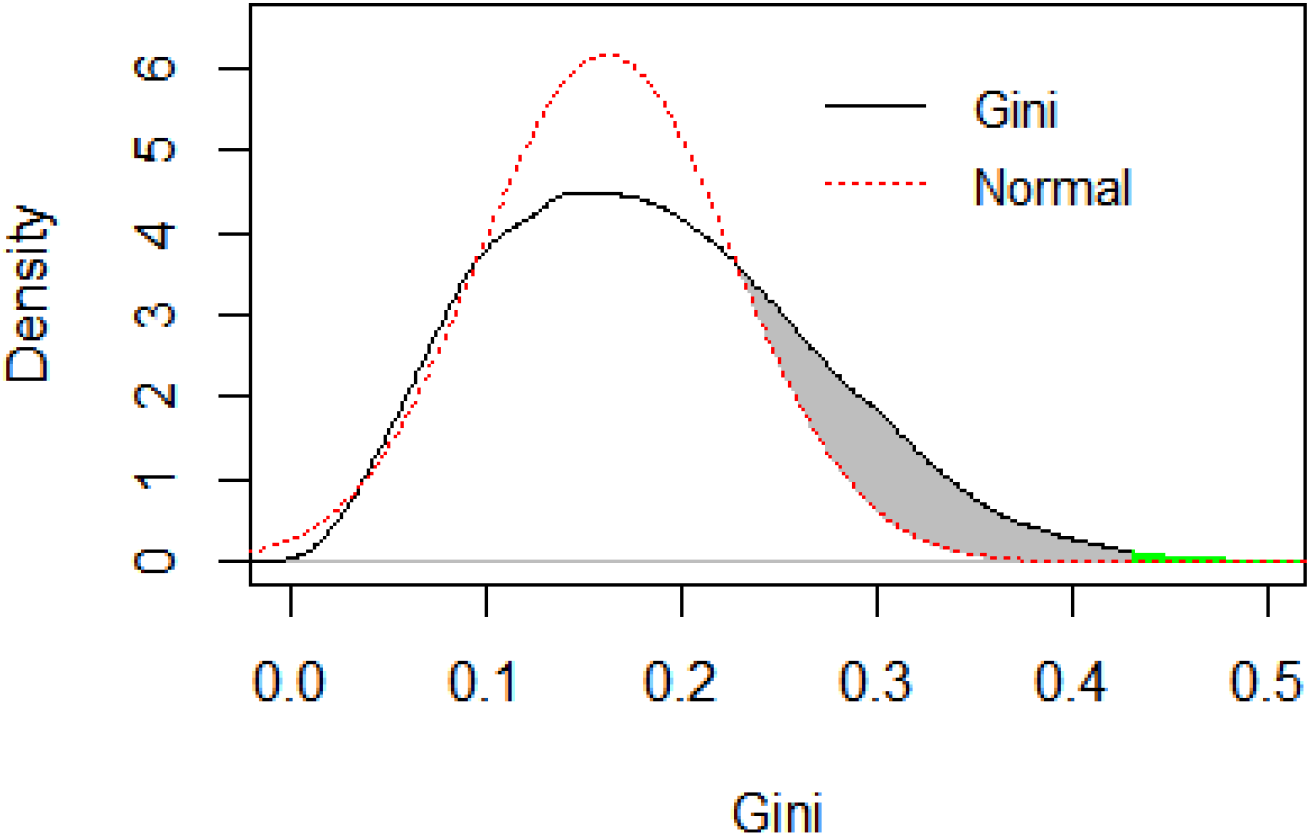
Gini index distribution (black curve) against its expected Gaussian counterpart (red curve). The gray area reflects the estimated excess of large Gini values. The green area marks the 81 extreme Gini values (FDR<0.05).

### Excessive self-citation suggests an author may be more likely to coerce others to cite their work

Because we were somewhat surprised to see %SC ranking correlating so well with all the NSC-based metrics (**Table 2**), we examined whether or not %SC effectively represents a risk factor for RLM. A prior small-scale study of Norwegian authors found a correlation between SC and NSC [23]. They hypothesized the increase in NSC was due to an “advertising” effect caused by SC. However, an alternative hypothesis might be that authors that place a high value on citation of their work might be inclined to use all venues available to them, citing themselves and asking others to as well.

**Figure 11** shows that, as an author’s %SC rises, their NSC-based Gini index FDR value drops. This means the more of their reference list an author reserves for self-citation, the more distorted their single-paper NSC frequency distribution is. Interestingly, the figure shows the average FDR curve flattening around 20% SC, suggesting that the ability of %SC to predict coercive NSC behavior has reached a point of diminishing returns. Recursive partitioning identified ≥18% SC as the optimal threshold for the group, separating a set of 948 authors at almost 50% risk (average FDR<=0.5), significantly higher than the rate for the group as a whole (t-test statistic=37, p<1e-16).

**Figure 11:**
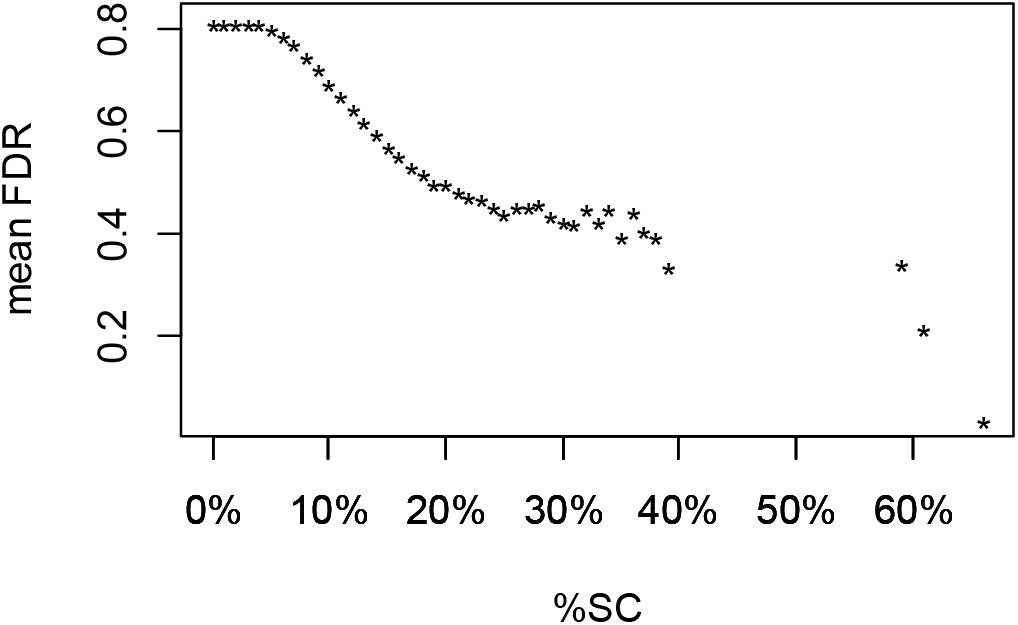
High %SC is correlated with signs of coercive behavior. For every author within each percentile of self-citation (x-axis), the mean Gini FDR (False Discovery Rate) is calculated for all authors above that percentile. Low Gini FDR values correspond to highly abnormal/skewed NSC patterns. If %SC were not predictive of NSC abnormalities, then the line would be approximately flat. Instead, the trend shows as %SC increases, authors greater than or equal to that amount have increasingly low Gini FDR values.

### Case Studies

We provide a list of all authors analyzed, their Gini values and their red flag metrics in **Supplementary Table 2**, which is available upon request. It is important to note that numbers are calculated using the PMC citation network subset and will be different from the same calculations (e.g., total papers & citations) derived using different sources such as ISI or Google Scholar. Our goal in this section is not to judge guilt or innocence, but to illustrate how a high Gini is associated with other unusual patterns (“red flags”) suggestive of citation hacking and how the red flag metrics lend themselves towards reasonable hypotheses regarding the potential origin of the distortion. We offer one example each of patterns suggestive of reviewer coercion, editorial coercion/influence, and author-author co-citation patterns that suggest mutual benefit.

The author with the highest Gini (Gini Rank #1), received 17+ NSC to his own papers 73 times, despite being in the 40^th^ percentile in total NSC, and has the highest observed to expected 17+ NSC ratio (114) by far among all authors. His MCJ index is also the highest among all authors (2.87), which is due to 72 of these 73 papers all coming from one journal (*Surgical Endoscopy*). Examining a random subset of these papers, we find they are predominantly from commentaries on other papers published in the journal rather than research papers. He has the highest rank in multiple red flag categories. He has the highest ratio of consecutive blocks per NSC (0.664), and the presence of very large blocks (>20) of consecutive citations were confirmed by manual examination of a random subset. He fell below threshold for fractional self-citation calculations with only 8 anchor author papers in the citation network, but averaged 32% self-citations in these 8. Interestingly, unusual self-citation and co-citation patterns for this author were reported in a prior study [9], which hypothesized that such a pattern suggested he was attempting to raise his H-index. Combined, this is highly suggestive of editorial citation hacking.

The author with Gini Rank #2, ranks in the 12^th^ percentile for total NSC, but received 17+ NSC to his work from 60 separate papers (Obs/Exp=42), the 3^rd^ highest. An estimated 35% of the extracted references in the papers he authored are self-citations (rank=19^th^ out of 20,803). His MCJI suggests these distortions, in general, are not attributable to influence at one specific journal. He has the 7^th^ highest number of blocks/NSC. Examining the context of the block citations in the published papers, we found a high degree of textual similarity surrounding them and that the context of the citations appears trivial (e.g., mentioning that user-friendly webservers are important followed by a very large number of citations to his papers). We also noticed that his name is mentioned frequently in the title of papers with excessive NSC, and that these have become increasingly frequent in recent years. Querying MEDLINE directly, we estimated almost 200 publications mention him by name in the title, but found only ~10% within our extracted citation network, suggesting the magnitude of his Gini distortion may be a significant underestimate. Googling textual phrases preceding large citation blocks to his papers (e.g., “to develop a really useful predictor”) shows that the same phrases appear in hundreds of papers verbatim as well as in a post-publication peer-review report online [24]. In this report, the reviewer requests 147 citations, the vast majority to this author, and after the first round of revision, rejects the paper because the authors did not accept the 1^st^-round request to change their title to include his name. This pattern is highly suggestive of reviewer-coerced citation, at least as a primary mechanism. However, we did observe an unusually large, but transient, surge of extreme NSC per paper coupled with his name being mentioned in the title of papers within two journals (*Prot Pept Let* from 2012-3 and *J Theor Biol* from 2018-9). We contacted *J Theor Biol* in 2019 because his activity was ongoing at the time, and an investigation by the editors in chief revealed that he had committed a number of ethical breaches as editor, including coercive requests to cite a large number of his papers [18].

The authors with Gini rank #3 and #8 cite each other extensively in papers they do not co-author, with #8 being the Most Citing Author (MCA) of #3 but not vice-versa. Having the same last name suggests they are related, the bylines in their papers shows they work at the same institution, the titles of their papers suggest they research the same subject, and PubMed shows they co-author frequently together. One of the reasons they may score so prominently is they also have a high rate of self-citation. A total of 19% and 20%, respectively, of their reference lists were self-citations, which puts them each in the top 3%. So on one hand, their NSC distortion could be attributed to a proclivity for self-citation plus entangled research activity but, on the other, it can be argued they each benefit from these co-citation patterns.

## Discussion

Some of the limitations of this study should be noted. First, the citation network we have is only a subset of the whole. References to papers will be biased towards the types of full-text articles that are most likely to be deposited in PubMed Central. Since the United States government mandates that publications resulting from federally funded research be deposited there, it will likely be biased towards authors in the US. Studies originating in and funded by other countries will likely be under-represented in the PMC citation network. Second, although our heuristics for author name disambiguation work reasonably well in general, they are imperfect and it biases our author subset towards authors with less ambiguous names. Consequently, authors with a national origin in which names are more ambiguous would be under-represented in this subset, so drawing demographic conclusions would be problematic. Third, the order in which citations appear in the XML files versus the PDFs is not preserved with high fidelity, so our calculations will underestimate the actual frequency and size of contiguous citation blocks to each author.

Prior to this study, citation hacking had only been studied via surveys and anecdotal reports. The number of currently active authors that we can declare, with 95% confidence, have significantly distorted NSC distributions was less than 1% of the total. Although this suggests the fraction of authors engaged in chronic citation hacking is relatively small, they have also been highly active in RLM and the percent of authors who have been hacked is much higher. Furthermore, the FDR suggests approximately 16% of authors have engaged in RLM to some degree at some point in the past.

Our analysis highlights the importance of having some system in place to detect and prevent citation hacking efforts before they become part of the published record. Once they do, it is unclear what should be done with compromised papers. In confirmed cases, removing the coerced references seems like a reasonable solution, but that may not be logistically simple and authors may be reluctant to cooperate if they feel admitting coercion might either tarnish their paper or possibly even their reputation. Alternatively, a database of reviewer/editor-added citations could be created, which could then be subtracted out of calculated metrics such as the H-index.

## Conclusion

If citations are the currency of science, then citation hacking is akin to counterfeiting. Trust in a currency is integral to its value and continued use. If we, as a scientific community, plan to continue using citations as a metric that reflects influence, then we need to be able to detect when the system may have been gamed. There are advantages to having a decentralized scientific publishing system, but one disadvantage is an inability to share information about bad actors within the system. Similarly, anonymous peer-review enables honest assessment without fear of repercussions, but also cloaks self-interested behavior. We cannot rely upon anecdotal reports coming from journals to identify bad actors, particularly if they are reluctant to name them [10, 18], as our data has shown cases of potential concern that started long ago and are still ongoing as of this writing. We’ve attempted to find the best solution within our grasp: A straightforward method to detect and prioritize; an unbiased study of all highly active, well-published authors; and the ability to put single instances of potential misconduct within a context. Along these lines we caution that, although it is almost inevitable that the efforts of citation hackers will leave “fingerprints” within the citation record, it does not necessarily follow that all authors with a high NSC Gini Index obtained them through unethical or coercive means.

## Acknowledgements

I would like to thank Alex Bateman, Lars Juhl Jensen and Hunter Porter for their valuable feedback on early versions of this manuscript.

## References

1. Lopez-Cozar, E.D., N. Robinson-Garcia, and D. Torres-Salinas, Manipulating Google Scholar Citations and Google Scholar Metrics: simple, easy and tempting. arXiv.org, 2012: p. 1212.0638.

2. Fong, E.A. and A.W. Wilhite, Authorship and citation manipulation in academic research. PLoS One, 2017. 12(12): p. e0187394.

3. Thombs, B.D., et al., Potentially coercive self-citation by peer reviewers: a cross-sectional study. J Psychosom Res, 2015. 78(1): p. 1–6.

4. Wilhite, A.W. and E.A. Fong, Scientific publications. Coercive citation in academic publishing. Science, 2012. 335(6068): p. 542–3.

5. Huggett, S., Journal bibliometrics indicators and citation ethics: a discussion of current issues. Atherosclerosis, 2013. 230(2): p. 275–7.

6. Biagioli, M., Watch out for cheats in citation game. Nature, 2016. 535(7611): p. 201.

7. Chen, C., et al., Emerging trends in regenerative medicine: a scientometric analysis in CiteSpace. Expert Opin Biol Ther, 2012. 12(5): p. 593–608.

8. Heneberg, P., From Excessive Journal Self-Cites to Citation Stacking: Analysis of Journal Self-Citation Kinetics in Search for Journals, Which Boost Their Scientometric Indicators. PLoS One, 2016. 11(4): p. e0153730.

9. Ioannidis, J.P., A generalized view of self-citation: direct, co-author, collaborative, and coercive induced self-citation. J Psychosom Res, 2015. 78(1): p. 7–11.

10. Wren, J.D., A. Valencia, and J. Kelso, Reviewer-coerced citation: case report, update on journal policy and suggestions for future prevention. Bioinformatics, 2019. 35(18): p. 3217–3218.

11. Martin, B.R., Whither research integrity? Plagiarism, self-plagiarism, and coercive citation in the age of research assessment. Res Policy, 2013. 42: p. 1005–1014.

12. Resnik, D.B., C. Gutierrez-Ford, and S. Peddada, Perceptions of ethical problems with scientific journal peer review: an exploratory study. Sci Eng Ethics, 2008. 14(3): p. 305–10.

13. Levis, A.W., et al., Comparison of self-citation by peer reviewers in a journal with single-blind peer review versus a journal with open peer review. J Psychosom Res, 2015. 79(6): p. 561–5.

14. Wren, J.D., et al., The write position. A survey of perceived contributions to papers based on byline position and number of authors. EMBO Rep, 2007. 8(11): p. 988–91.

15. Ghosh, A., et al., Zipf’s law in city size from a resource utilization model. Phys Rev E Stat Nonlin Soft Matter Phys, 2014. 90(4): p. 042815.

16. Schwab, D.J., I. Nemenman, and P. Mehta, Zipf’s law and criticality in multivariate data without fine-tuning. Phys Rev Lett, 2014. 113(6): p. 068102.

17. Yong, A., Critique of Hirsch’s Citation Index: A Combinatorial Fermi Problem. Notices of the American Mathematical Society, 2014. 61(11): p. 1040–50.

18. Chaplain, M., D. Kircshner, and I. Yoh, JTB Editorial Malpractice: A Case Report. Journal of Theoretical Biology, 2020. (in press).

19. Ioannidis, J.P.A., et al., A standardized citation metrics author database annotated for scientific field. PLoS Biol, 2019. 17(8): p. e3000384.

20. Van Noorden, R. and D. Singh Chawla, Hundreds of extreme self-citing scientists revealed in new database. Nature, 2019. 572(7771): p. 578–579.

21. Lubrano, M., The econometrics of inequality and poverty. Lecture 4: Lorenz curves, the Gini coefficient and parametric distributions. 2013.

22. Ultsch, A. and J. Lotsch, A data science based standardized Gini index as a Lorenz dominance preserving measure of the inequality of distributions. PLoS One, 2017. 12(8): p. e0181572.

23. Fowler, J.H. and D.W. Aksnes, Does self-citation pay? Scientometrics, 2007. 72(3): p. 427–437.

24. ; Available from: https://www.mdpi.com/1422-0067/21/1/75/review_report.

